# exFINDER: identify external communication signals using single-cell transcriptomics data

**DOI:** 10.1101/2023.03.24.533888

**Authors:** Changhan He, Peijie Zhou, Qing Nie

**Affiliations:** Department of Mathematics, University of California, Irvine, Irvine, CA 92697, USA; Department of Cell and Developmental Biology, University of California, Irvine, Irvine, CA 92697, USA

## Abstract

Cells make decisions through their communication with other cells and receiving signals from their environment. Using single-cell transcriptomics, computational tools have been developed to infer cell-cell communication through ligands and receptors. However, the existing methods only deal with signals sent by the measured cells in the data, the received signals from the external system are missing in the inference. Here, we present exFINDER, a method that identifies such external signals received by the cells in the single-cell transcriptomics datasets by utilizing the prior knowledge of signaling pathways. In particular, exFINDER can uncover external signals that activate the given target genes, infer the external signal-target signaling network (exSigNet), and perform quantitative analysis on exSigNets. The applications of exFINDER to scRNA-seq datasets from different species demonstrate the accuracy and robustness of identifying external signals, revealing critical transition-related signaling activities, inferring critical external signals and targets, clustering signal-target paths, and evaluating relevant biological events. Overall, exFINDER can be applied to scRNA-seq data to reveal the external signal-associated activities and maybe novel cells that send such signals.

## INTRODUCTION

The transformation from a single cell to a multicellular organism is built upon the development of individual cells, including cell proliferation (1), differentiation (2), migration (3), and death (4). Numerous works have revealed cell decision is regulated by microenvironmental signals (5, 6). Understanding the cell fate decisions has been a major challenge (7-10). Recent advance in single-cell transcriptomic data (scRNA-seq) allows unprecedented resolution in identifying the diversity of cellular states and uncovering signaling activities in cell lineage decision (11-13).

Cell-cell communication is critical to cell fate decisions and many other biological processes (14-16). Recently, computational methods with different methodologies and functionalities have been developed in identifying and quantifying cell-cell communication using scRNA-seq data. For example, SoptSC (17) infers communication between individual cells; while many other methods such as COMUNE (18), SingleCellSignalR (19), and CellChat (20), infer the cell-cell communication between cell clusters. In modeling ligand-receptor interactions, CellPhoneDB (21) and ICELLNET (22) take the multi-subunit structure of ligands and receptors into account, and CellChat (20) considers not only such structure but also the impact of agonists and antagonists. For downstream analysis, NicheNet (23) and scMLnet (24) infer multilayer signaling networks linking ligands and target genes. Those methods have been applied to many different systems, including diseases, to find novel signaling molecules (25-27), and their different functionalities have been compared and studied (28, 29).

Besides the communication among cells measured in the collected datasets, the communication between the measured cells and the external system, which may include the unmeasured cells (30), extracellular signals (31, 32), or induced signals (33), plays important roles in the functions of those measured cells. For example, the signals (ligands) from the environment, which are not expressed but received by the measured cells, induce the epithelial-mesenchymal transition (EMT) (34), and result in migration, invasion, EMT of prostate cancer cells (35). However, when inferring cell-cell communication, the current methods only deal with ligand-regulated signals sent by the measured cells in the data; that is, the signal must be related to a ligand highly expressed in the measured cells. As a result, those methods fail to consider the received ligands from the external system nor the non-cellular components that can hardly be directly measured using scRNA-seq. For example, a recent study (34) has shown that inducers *TGFB1*, *EGF*, and *TNF*, not expressed in the measured cells, can all serve as promotors during the epithelial-mesenchymal transition (EMT); and it has been found that *Gdf9* increases the reprogramming efficiency and the fraction of cells with neural fates (33). Identifying such external signals and analyzing the corresponding signaling pathways are critical to revealing important factors in cell fate transitions (34,36,37) and disease (35,38,39).

Here we develop exFINDER, a method and an open-source R package to (a) identify external signals that activate target genes; (b) infer the signaling pathways linking external signals and target genes; and (c) quantitatively analyze the network. Specifically, we first collect and integrate multiple complementary data sources containing ligand-to-target signaling paths based on prior knowledge, and then infer the ligand-target Gene Regulatory Network (GRN) starting from the given target genes. Next, based on the inferred ligand-target GRN, exFINDER identifies the external signals and infers the external signal-target signaling network (exSigNet) within a given scRNA-seq data using mass action models. Moreover, exFINDER quantitatively analyzes the exSigNet by predicting signaling strength, calculating the maximal signal flow, clustering different ligand-target signaling paths, quantifying the signaling activities using the activation index (AcI), and evaluating the GO analysis outputs of exSigNet. In addition, exFINDER provides several intuitive visualization outputs to interpret exSigNet and quantitative analysis outputs. We demonstrate the accuracy and robustness of exFINDER by applying the method to publicly deposited scRNA-seq datasets of different species, including human, zebrafish, and mouse.

## MATERIALS AND METHODS

The method exFINDER performs three major functions: 1) inferring ligand-target GRN from the given target genes based on prior knowledge using exFINDER database (exFINDER-DB), 2) identifying external signals and inferring the external signal-target signaling network (exSigNet) from the ligand-target GRN using scRNA-seq data, 3) quantitatively analyzing the exSigNet on its various properties and visualizing the outputs.

### Database integration and inference of ligand-target GRN based on prior knowledge

A database of signaling pathways from ligands to target genes is required to find what ligands activate specific target genes. Here we collect and integrate multiple complementary data sources to obtain such a database and use it to infer the ligand-target GRN.

#### Database integration based on publicly available sources

To obtain a signaling network database that comprehensively represents the signaling from ligands to target genes, we consider three layers of information: ligand-receptor interactions, signal transductions from receptor to transcription factors (TFs), and transcription factor-target regulatory interactions. For exFINDER database (exFINDER-DB), we integrate several publicly available databases to augment the NicheNet database (23) by including 1) the CellChat database (20) and the IUPHAR database (40) on ligand-receptor interactions, 2) the Garcia-Alonso database (41) on transcription factor-target regulatory interactions. Furthermore, for each signaling path, we label the sender-receiver pair based on their gene symbols and label the corresponding sources. The exFINDER-DB is made available for human, mouse, and zebrafish. Because exFINDER-DB mines and combines multiple databases (Table S1), naturally it contains more interactions with more different species.

#### Inference of ligand-target GRN from the given target genes

We first select a set of genes (denoted as *V_T_*) as the target genes. The target genes in performing exFINDER analysis need to be part of the measured genes in the data, and they are selected based on prior knowledge (e.g., other experimental data), prediction (e.g., differentially expressed genes), or genes of the user’s interest. Specifically, we apply exFINDER to infer the transcription factors (*V_TF_*) that regulate the target genes as well as the receptors (*V_R_*) interacting with these transcription factors. Finally, we infer the ligands (*V_L_*) that interact with the receptors. All these inferences are obtained using the exFINDER-DB. Moreover, for interaction from 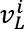 (the *i^th^* ligand in *V_L_*) to 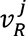, we denote it as 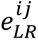, and define *E_LR_* as the set of all inferred ligand-receptor interactions. Similarly, we define *E_RTF_*, and *E_TFT_*. Next, we convert these genes and the signaling paths between them into a directed graph *G = 〈V, E〉*, where *V* = *V_L_* ∪ *V_R_* ∪ *V_TF_* ∪ *V_T_* is the node set and *E* = *E_LR_* ∪ *E_RTF_* ∪ *E_TFT_* is the edge set. This directed graph *G* is named as the ligand-target GRN, which is inferred by only using prior knowledge.

### Identification of external signals and inference of the external signal-target signaling network (exSigNet) from the ligand-target GRN using scRNA-seq data

Once we have the ligand-target GRN and the scRNA-seq data as inputs, exFINDER can identify the external signals and infer the signaling paths from the external signals to target genes using the scRNA-seq data, which are denoted as the exSigNet. Such analysis takes three steps:

#### Calculation of the average expression levels for all ligands

First, we calculate the average expression level of every ligand in each cell group. We first calculate the average expression levels of each gene in every cell population group. For every cell population group, we remove the genes with zero average expression and then calculate the 50^th^ percentile (42, 43) and the 90^th^ percentile (44, 45) expression levels. Next, the cutoff levels for “lowly expressed” genes are chosen to be below the maximal 50^th^ percentile among all cell groups, and the cutoff levels for “highly expressed” genes are chosen to be above the minimal 90^th^ percentile of all cell groups. This method takes into consideration the expression variation across cell populations as well as cutoff values with very large differences, providing a natural way to select the “lowly expressed” and “highly expressed” genes. Next, we mark those lowly expressed ligands among all cell groups as potential external signals.

#### Identification of external signals

Within the ligand-target GRN, we first select highly expressed transcription factors and their linked highly expressed receptors, which indicate possible signaling activities. For the potential external signals that interact with these receptors, their signaling pathways are inferred based on prior knowledge, and their downstream genes are highly expressed, which ensures they are external signals.

#### Inference of the exSigNet

Based on the above analysis, we connect the signal pathways from the external signals to the target genes, and then we convert such network into a directed graph *Ĝ* = 〈*V̂*, *Ê*〉, where *V̂* is the set of nodes and *Ê* is the set of edges. The exSigNet is denoted by *Ĝ*, which is a subgraph of *G*.

### Quantitative analysis of the exSigNets using scRNA-seq data

To better quantify the signaling activities and understand the functions of the exSigNets, exFINDER predicts the signaling strength and provides several quantitative analysis options.

#### a) Prediction of signaling strength within the exSigNet

First, we calculate the average expression levels of the receptors, transcription factors, and target genes within the exSigNet in each cell group. Second, since signaling activities may not occur in all measured cells, we select these genes’ maximal average expression levels among all cell groups to predict the signaling strength. Meanwhile, besides using the maximal average expression level of all cell groups, users can manually assign or define specific cell groups in exFINDER if needed. For external signals, we set their expression levels to one to avoid drastically affecting the prediction. Furthermore, if users want to check the external signals’ expression in specific cell groups, which may be measured in other scRNA-seq data, they can always load the corresponding data and assign the cell groups in exFINDER to calculate the external signals’ expression levels. To predict the signaling strength of an interaction, we use the mass action law based on the following model (20):

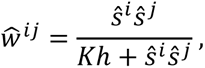

where *Ŝ^i^* and *Ŝ^j^* represent the expression levels of gene *v̂^i^* and *v̂^j^* in *Ĝ*, respectively. *ŵ^ij^* is the predicted signaling strength of the interaction between *v̂^i^* and *v̂^j^* (*ê^ij^*). Here *Kh* is the apparent dissociation constant (or half-saturation constant), meaning if *Ŝ^i^Ŝ^j^* = *Kh*, then *ŵ^ij^* = 0.5 reaches the half of the maximal strength. For simplicity we set its default value to two. By testing the model with different values of *Kh* between 1 to 10 using both synthetic and experimental data (46), we find although changing the *Kh* value might affect the absolute levels of the interactions, their relative signaling strengths remain hardly changed (Supplementary Figure S1).

#### b) Calculation of the maximal signal flow between single external signal-target pairs

After predicting the signaling strength of an exSigNet *Ĝ*, we next infer how the exSigNet connects a specific external signal-target pair. If such a network does not exist, then the maximal signal flow is zero. Otherwise, we use the Goldberg-Tarjan algorithm (47) to calculate the maximum signal flow from a given external signal to a given target in order to quantify the overall signaling strength in between. By comparing the sum of maximum signal flows from one external signal to all target genes (i.e., the total signal outflow), we predict the critical external signals in exSigNet *Ĝ*. Similarly, critical target genes are also predicted based on the total signal inflows.

#### c) Clustering of different external signal-target pairs within the exSigNet Ĝ

Since exSigNet *Ĝ* may include multiple external signals and target genes, quantifying the similarities of signaling activities between these external signal-target pairs helps analyze the functionalities of exSigNet *Ĝ*. Here, we first infer the exSigNets of every external signal-target pair from *Ĝ*, and then build a strength matrix *W* presenting the signaling strength of all the edges of *Ĝ*. The matrix *W* has (|*V̂_L_*| · |*V̂_T_*|) rows and |*Ê*| columns. Next, we normalize matrix *W* (for noise reduction) and perform *k*-means clustering for all external signal-target pairs.

#### d) Quantification of the signaling activity of the exSigNet Ĝ using the activation index (AcI)

To quantify the signaling activity of *Ĝ*, we use the activation index (AcI), considering both the signaling strength and the overall complexity of the network. The AcI of *Ĝ* is modeled by:

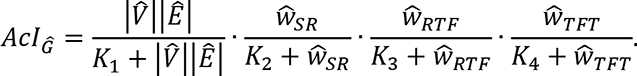

Here |*V̂* | and |*Ê*| is the total number of nodes and edges of *Ĝ*. *K_i_* (*i* = 1,2,3,4) are parameters with default value two. *ŵ_LR_*, *ŵ_RTF_*, and *ŵ_TFT_* represent the total signaling strength in three layers, respectively.

#### e) Evaluation of GO analysis outputs of exSigNet Ĝ using two quantities

To connect the exSigNet *Ĝ* to the GO analysis outputs, exFINDER uses two quantities to measure the GO terms projected to *Ĝ*. For each GO term, we identify the GO term-related genes in exSigNet and denote them as *V_GO_*. Then we calculate the proportion of genes involved in this GO term by 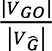, and compare the total expression level of these involved genes to the entire exSigNet using 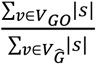, here *s* is the expression level of gene *v* calculated in part (a).

### Parameter explanation

In this study, when identifying the differentially expressed genes, we used the *FindMarkers* function in Seurat (v4.1.3) and followed the official workflow, using the Wilcoxon Rank Sum test with *min.pct = 0.25* and *logfc.threshold = 0.25*. When clustering the external signal-target pairs within the exSigNet, we used the *kmeans* function with 20 random sets and default “Hartigan-Wong” algorithm. When performing GO enrichment analysis, we used the *enrichGO* function in clusterProfiler (v4.2.2) with the default settings (*pvalueCutoff = 0.05, pAdjustMethod = “BH”, qvalueCutoff = 0.2, minGSSize = 10, maxGSSize = 500*). For critical transition analysis using BioTIP, we followed the official workflow with default settings. For determining the cutoff levels of lowly and highly expressed genes, we calculated the 50^th^, 75^th^, and 90^th^ percentiles for all datasets used in our study (Supplementary Figure S2). To facilitate better understanding of the method, we summarize the defined terminologies and concepts introduced in this study as a table (Table 1).

**Table 1.** A list of terminologies and concepts used in exFINDER.

| **Concept** | **Definition** | **References** |
| --- | --- | --- |
| external system | unmeasured cells, extracellular signals or induced signals that communicate with the measured cells | New |
| external signal | ligands that are received by measured cells but come from the external system | New |
| external signal-target signaling network (exSigNet) | signaling network that starts from external signals to the target genes | New |
| activation index (Acl) | index that quantifies the signaling activity of the exSigNet | New |
| maximal signal flow | derived from the “maximum flow problem”, which is defined as the maximal value of flow from the source (external signal) to the sink (target gene), and the signaling strength is regarded as the capacity of the edge | (80) |
| total signal outflow | the sum of maximum signal flows from one external signal to all target genes | New |
| total signal inflow | the sum of maximum signal flows from all external signals to one target gene | New |
| critical external signal | the external signal with the highest total signal outflow in the exSigNet | New |
| critical target gene | the target gene with the highest total signal inflow in the exSigNet | New |
| critical transition (CT) | process in which cells shift abruptly from one state to another | (81) |
| critical transition index (IC) | index used to predict the critical transition | (59) |
| critical transition signal (CTS) | a small group of genes identified by an existing method (BioTIP) to characterize regulated stochasticity during semi-stable transition | (60) |

## RESULTS

### Overview of exFINDER

For the inputs, exFINDER requires gene expression data from cells, user-assigned cell labels, and user-selected target genes that may be activated by the sought external signals (Figure 1A). For example, those genes can be critical genes in differentiation or maker genes for specified cell groups. Such information can be obtained using other computational methods such as Seurat (48-51) or CellChat (20). During the exFINDER analysis, users may also load additional cell groups that are specific for external signal identification. Other relevant analysis, such as marker genes identification, lineage trajectory and critical transition construction (Figure 1B), are useful when analyzing external signals associated with cell fate decision activities, such as differentiation. With the input data, exFINDER performs the tasks in the following modules:

**Figure 1.**
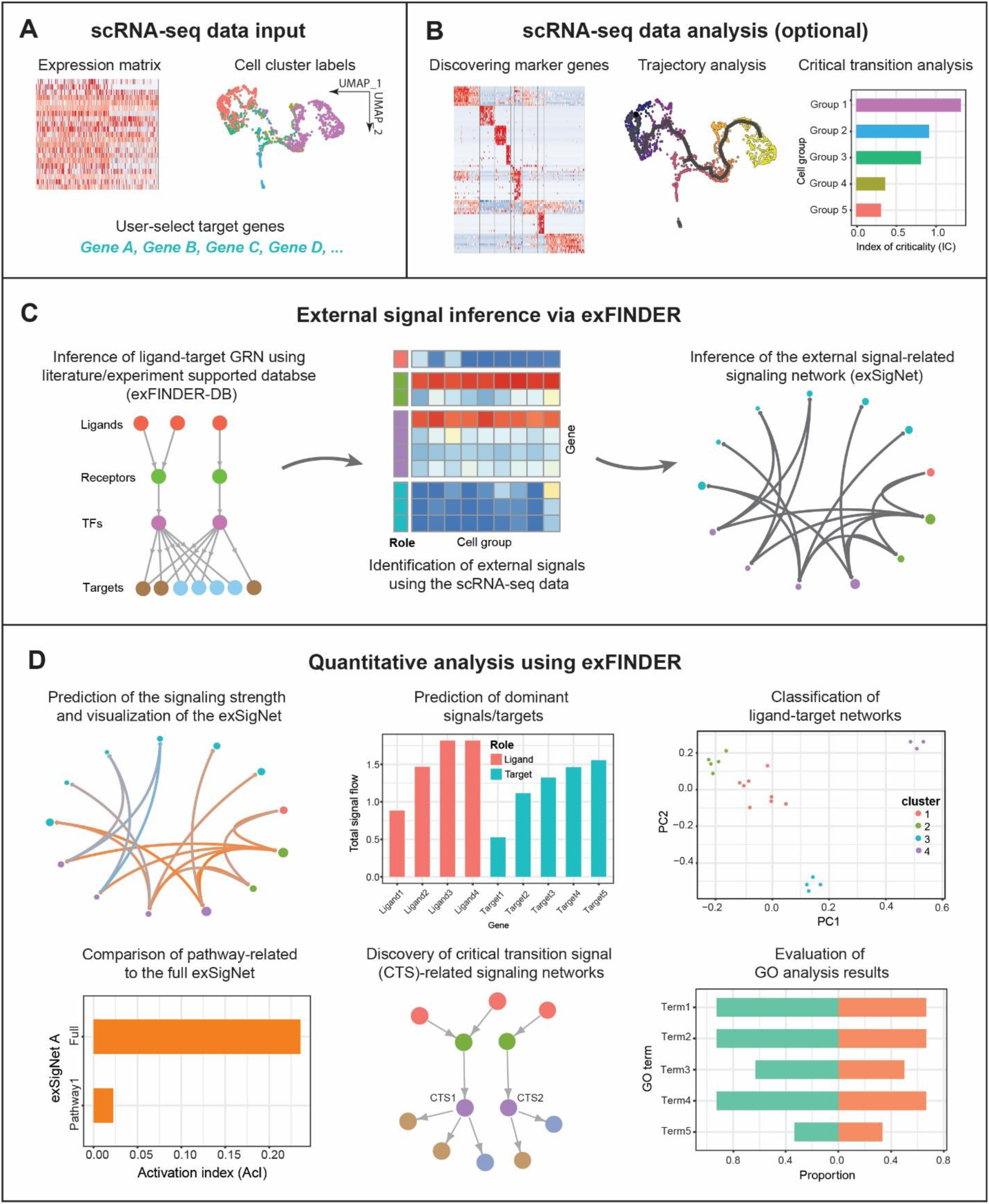
Overview of exFINDER. **(A)** exFINDER requires scRNA-seq data, user-assigned cell cluster labels and user-selected target genes as inputs. **(B)** Additional information, such as differentially expressed genes, pseudo time, and critical transition analysis information, can be included for specific biological problems. **(C)** exFINDER first infers the ligand-target GRN based on the structure of ligand-receptor-transcriptional factors-target (L-R-TF-T) from exFINDER-DB. It identifies external signals using the scRNA-seq data and infers their corresponding exSigNet. **(D)** exFINDER predicts the signaling strength, visualizes the exSigNet, and quantitatively analyzes the networks through graph theory methods for interpretation of exSigNet, including identifying critical ligands and target genes, classifying networks between different ligand-target pairs, finding external signal-activated pathways, uncovering critical transition-related signaling networks, and evaluating GO analysis outputs.

#### 1. Database integration and inference of ligand-target GRN from the given target genes

Here we link ligands and targets by considering the “L-R-TF-T” signaling structure. We integrated multiple data sources (see Methods) and obtained the exFINDER database (exFINDER-DB), which is available for human, mouse, and zebrafish. With the exFINDER-DB, we next employ exFINDER to infer all relevant transcription factors, receptors, and ligands with the “L-R-TF-T” signaling structure from the given target genes. Then exFINDER converts this network into a directed graph *G = 〈V, E*〉, where *V* is the set of nodes in *G* that represents the genes, and *E* is the set of edges and contains all the inferred signaling paths (Figure 1C, see Methods).

#### 2. Identification of external signals and inference of the external signal-target signaling network (exSigNet) from the ligand-target GRN using scRNA-seq data

exFINDER identifies the external signals based on their low or no expressions in the data and the high expression of their downstream genes. Then it infers the signaling network connecting external signals and target genes into a directed graph *Ĝ* = 〈*V̂*, *Ê*〉, which represents the exSigNet and is also a subgraph of *G* (Figure 1C, see Methods).

#### 3. Quantitative analysis of the exSigNet Ĝ and visualizations

To predict the signaling strength of each interaction (i.e., edge) in the exSigNet *Ĝ*, we calculate the expression level of the genes (i.e., nodes) in *Ĝ* and model the signaling strength via mass action (see Methods). exFINDER provides informative and intuitive visualizations via both customized circle plots and hive plots for the exSigNet (Figure 1C-D). To uncover the functionally related ligand-receptor interactions activated by *Ĝ*, we compare the external signal-receptor interactions to a published database (20). For the functionally related interactions activated by *Ĝ*, exFINDER infers their own exSigNets (Figure 1D) and compares their activation index (AcI) using a bar plot (see Methods). To reveal the critical transition-associated signaling, we next determine the roles of the critical transition signals (CTSs) in the “L-R-TF-T” signaling structure, a small group of transcription factors that regulate the transition of cell states (Table 1). Then we infer their corresponding ligand-target GRN based on *Ĝ* using a tree for visualization (Figure 1D). Furthermore, exFINDER performs a variety of analyses of *Ĝ* in an unsupervised manner. First, it predicts the critical external signals and target genes of *Ĝ* by their maxima signal outflows and inflows, respectively (see Methods); Second, for all external signal-target pairs, exFINDER infers their own exSigNets and uses clustering learning to showcase their similarities (Figure 1D); Third, exFINDER evaluates the exSigNet *Ĝ*’s GO analysis outputs by calculating the proportion of involved genes and their expression levels (Figure 1D, see Methods). Overall, these functionalities allow exFINDER to uncover external signals, reveal mechanisms in activating the target genes, and predict novel cell groups.

### Benchmarking exFINDER using subsets of cells measured in the datasets for human and mouse

To benchmark exFINDER, we used subsets of cells in a dataset to infer ligand-receptor communication receiving from the rest cells in the dataset. Specifically, we evaluated its performance on using a subset of the published human skin scRNA-seq dataset (52) containing four cell groups: two subpopulations of dendritic cells (cDC2 and LC) and two subpopulations of fibroblasts cells (*FBN1*^+^ FIB and *APOE*^+^ FIB). We first found differentially expressed genes of each cell group (Supplementary Figure S3A), then performed CellChat analysis to infer cell-cell communication between these cell groups (Figure 2A-B, Supplementary Figure S3C-D) and exported the ligands targeting each cell group.

**Figure 2.**
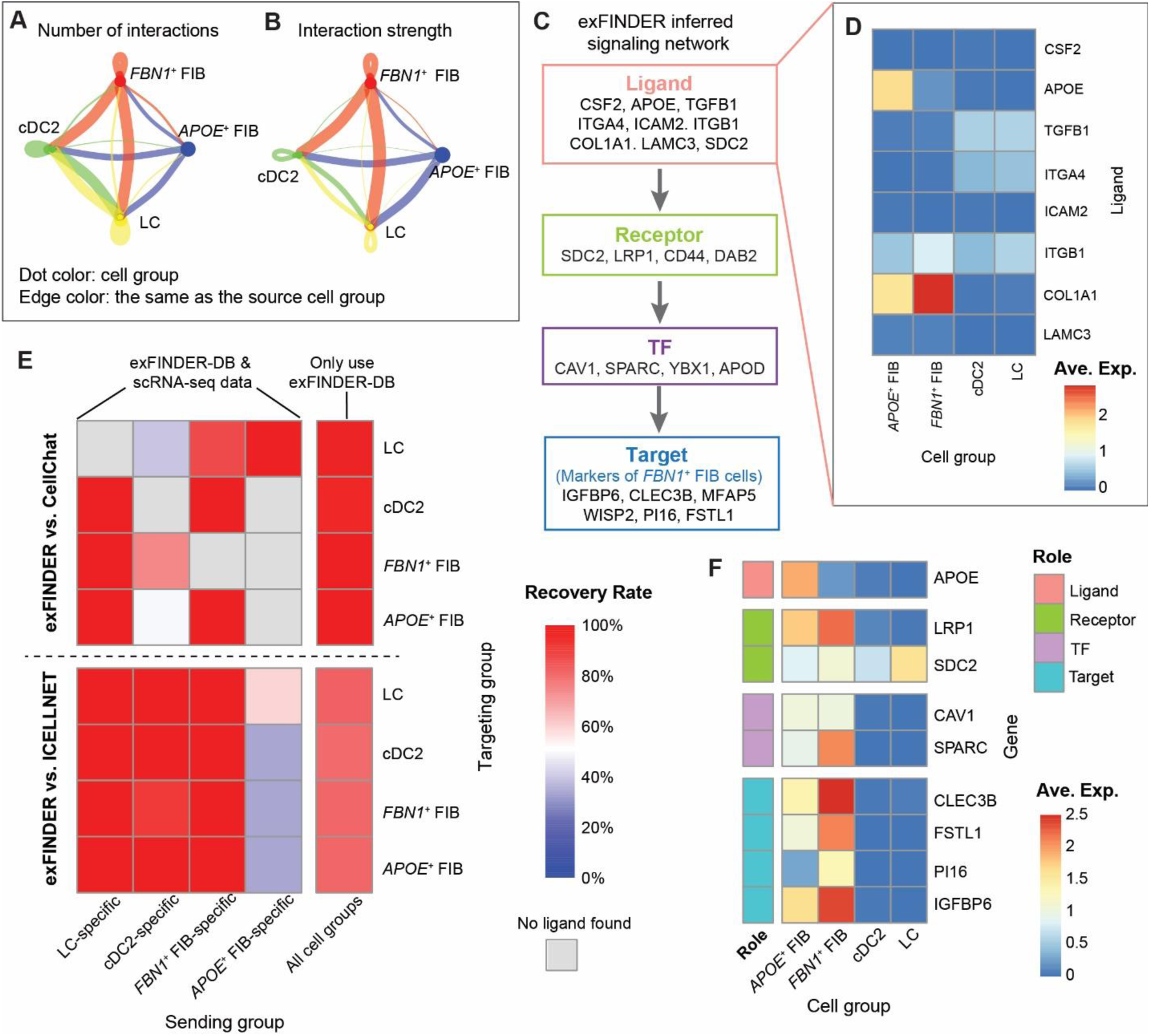
Benchmarking exFINDER using human skin data. **(A-B)** The significant ligand-receptor interactions among four cell populations inferred by CellChat. Each dot color represents one cell group, and the edge color is the same as the cell group sending the signal. The dot size is proportional to the population size of the indicated cell group. The edge width is proportional to the indicated number (A) and strength (B) of ligand-receptor pairs. **(C)** A schematic of the exFINDER-inferred signaling network targeting *FBN1*^+^ FIB cells. **(D)** Heatmap of the expression level of all inferred signals across four cell population groups, with ligand *APOE* identified as an external signal. **(E)** Heatmap showing the percentages of ligands inferred by CellChat and ICELLNET as well as captured by exFINDER. The X-axis represents the cell population groups expressing the ligands, and Y-axis represents the cell population groups receiving the signals. The color bar is the percentages of CellChat (top) and ICELLNET (bottom)-inferred ligands that are also identified by exFINDER. The 5^th^ column shows the ligand recovery rate of exFINDER only using prior knowledge, and the rest four columns show the ligand recovery rate of exFINDER using both prior knowledge and the scRNA-seq data. And the grey block indicates no such ligands inferred by CellChat or ICELLNET. **(F)** Heatmap of the expression level of the components in the exSigNet built by exFINDER.

For each cell group, we employed exFINDER to infer the ligand-target GRN targeting its top 10 marker genes using prior knowledge and found the ligands targeting each cell group. For example, six of the top 10 marker genes of *FBN1*^+^ FIB cells are regulated by nine ligands, including only one highly expressed in *APOE*^+^ FIB cells (*APOE*), one comes from both *APOE*^+^ FIB and *FBN1*^+^ FIB cells (*COL1A1*), while others come from the external environment (Figure 2C-D). Although exFINDER might infer more ligands than CellChat, including the ones that are lowly expressed or even unmeasured, we only compared them to the four groups of ligands inferred by CellChat to check how many of them were captured by exFINDER. In this way, we study its capability in inferring ligands produced by the measured cells. We found that exFINDER successfully recovers all CellChat-inferred ligands only using the exFINDER-DB (Figure 2E, the 5^th^ column), suggesting good coverage and robustness of exFINDER-DB.

Then we tested exFINDER’s capability of identifying external signals. Each cell group was removed from the data such that they became the “external cells” (i.e., taking the cells out of the data to make them as unmeasured cells) for the rest cells in the data. Obviously, CellChat is unable to infer the ligands produced by such “external cells”. Taking the *APOE*^+^ FIB cells as an example, *APOE* is an *APOE*^+^ FIB cell-specific ligand (i.e., only highly expressed in the *APOE*^+^ FIB cells) that targets the *FBN1*^+^ FIB cells (Figure 2D). By setting the *APOE*^+^ FIB cells to be the “external cells”, CellChat couldn’t find *APOE* as a ligand. On the other hand, exFINDER successfully identified *APOE* as an external signal and inferred the exSigNet (Figure 2F).

Similar analyses were also performed on the other cell groups. For the cell type-specific ligands, CellChat failed if the corresponding cell group was missing in the data, while exFINDER could always identify them as external signals. In most cases (8 out of 10), exFINDER is able to recover over 80%, with 40% in the worst case (Figure 2E).

Meanwhile, we also compared the performance on inferring ligands and external signals of exFINDER and ICELLNET, a computational method to infer ligand-receptor interactions (22). We first performed ICELLNET analysis and selected the ligand-receptor pairs with positive communication probability. For the ligands inferred by ICELLNET targeting different cell groups, exFINDER always captures over 80% of them by only using the prior knowledge (Figure 2E, the 5^th^ column). And in most cases (12 out of 16), for the ligands that ICELLNET cannot capture (when the corresponding source cell group was removed), exFINDER is able to recover over 90%, with 33% in the worst case (Figure 2E).

Moreover, a similar comparison between exFINDER and CellChat was carried out by using embryonic mouse skin scRNA-seq dataset (53). We first selected a subset of cells containing four cell groups (Immune, ENDO, MYL, and MELA), then performed CellChat analysis (Supplementary Figure S3B, S3E-F) and compared with the exFINDER results. We found that over 85% recovery rate in capturing CellChat-inferred ligands, and over 65% recovery rate in identifying external signals in most cases (7 out of 10) (Supplementary Figure S4).

In fact, both CellChat and ICELLNET infer communication links only based on the expression levels of the ligands and receptors whereas exFINDER ensures the ligand-related signaling pathway eventually activates the downstream marker genes. Such comparison partially shows the coverage and robustness of exFINDER-DB and good accuracy in identifying external signals by exFINDER.

### exFINDER identifies differentiation-associated external signals during zebrafish neural crest (NC) development

To study exFINDER’s ability in recovering external signals that affect cell differentiation, we analyze a scRNA-seq dataset for the cranial NC cells that contribute to zebrafish’s first pharyngeal arch (PA1) (46). A previous study shows cells differentiate from early NC cells to pigment cells and skeletal cells with the presence of transitional cells during this differentiation process occurring at around 18 hpf (Figure 3A, Supplementary Figure S5A-C). In this study, only the NC cells (six subgroups) were analyzed without including the non-NC cells (four subgroups) that were measured (46) (Supplementary Figure S5D). Our CellChat analysis on all cells suggests strong communications between NC cells and non-NC cells (Figure 3B-C), indicating that NC cells may have received external signals produced by the non-NC cells. Meanwhile, signals from the external environment (i.e., not expressed in the measured cells) may also be received to affect the differentiation.

**Figure 3.**
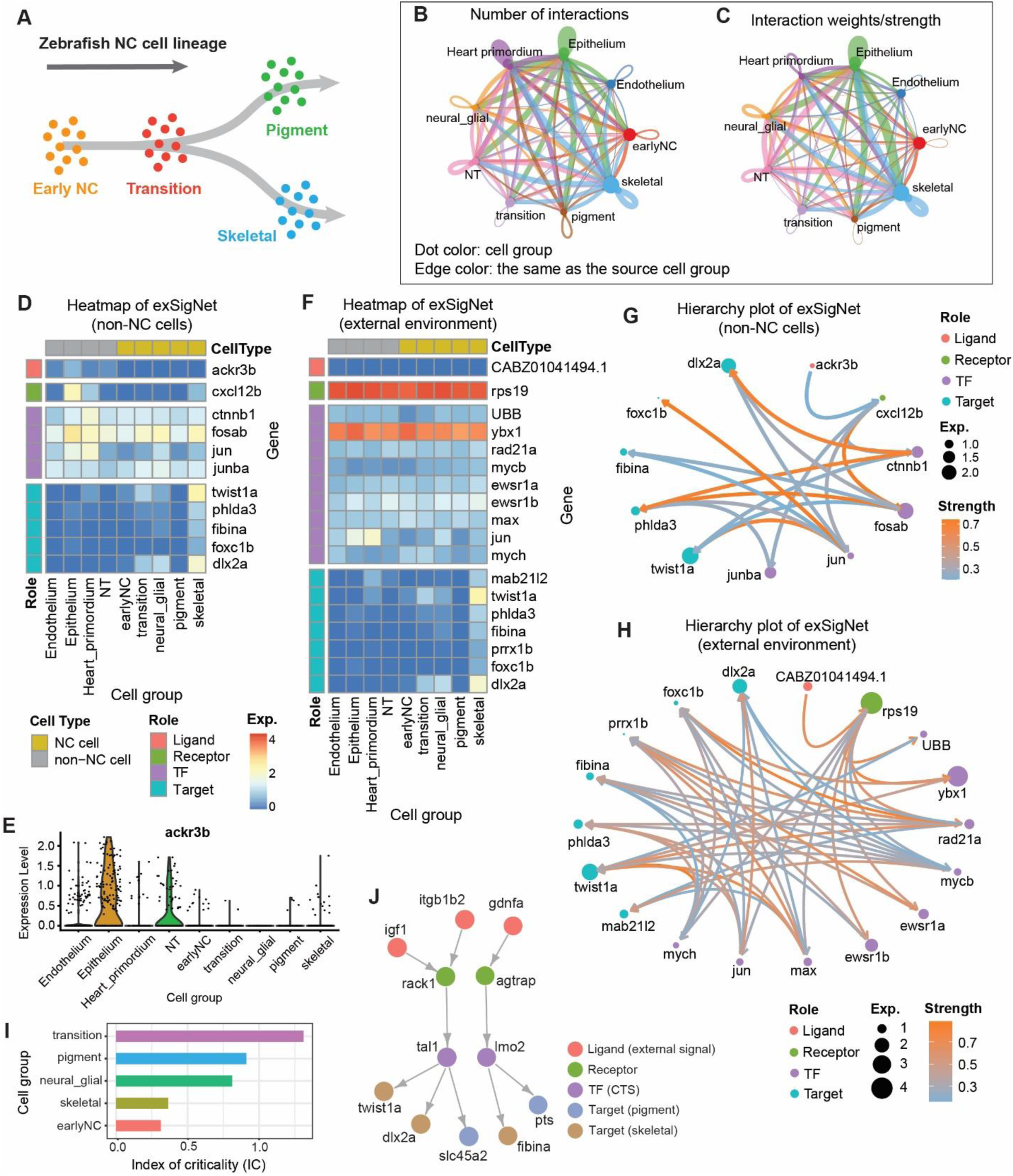
exFINDER identifies cell differentiation-associated external signals during zebrafish neural crest (NC) development. **(A)** Schematic of neural crest cell lineage, showing the cell differentiation process along the timeline (46). **(B-C)** CellChat analysis for the number and strength of ligand-receptor interactions between different cell populations. **(D)** Expression heatmap of the exSigNet associated with the inferred signals produced by non-NC cells targeting skeletal cell marker genes. **(E)** Expression levels of inferred signal *ackr3b* in each cell group. **(F)** Expression heatmap of the exSigNet associated with the inferred signals coming from the external environment targeting skeletal cell marker genes. **(G-H)** Circle plots showing the exSigNets. Node size is proportional to the gene expression level, and the color bar represents the predicted signaling strength. **(I)** Index of criticality of different NC cell groups generated by BioTIP. **(J)** Critical transition signal-involved exSigNet inferred by exFINDER.

To find the external signals produced by non-NC cells or the external environment that may drive differentiation, we first identified the marker genes of all cell groups and inferred the ligand-target GRN from the top 10 marker genes of skeletal cells (Supplementary Figure S5E-F). Based on this ligand-target GRN, exFINDER found that *ackr3b* is only highly expressed in the non-NC cells, and linked to five targeted marker genes (*twist1a*, *phlda3*, *fibina*, *foxc1b* and *dlx2a*) (Figure 3D-E). The external signal *ackr3b* activates the target genes through highly expressed downstream genes (Figure 3D). These findings are consistent with previous studies showing that *ackr3b* is expressed in a wide range of tissues during somitogenesis, including central nervous system and somites (54), along with recent evidence on the interactions between *ackr3b* and *cxcl12b* (55). Furthermore, signal *CABZ01041494.1* (also denoted as *c5ar1* based on its Ensembl: ENSDARG00000040319) was found from the external environment as seen in its exSigNet (Figure 3F). Based on the exFINDER-DB, its receptors *rps19*, which is connected with seven marker genes through nine TFs, is only activated by *CABZ01041494.1*. This result is consistent with a previous study showing the interactions between *C5aR1* and *RPS19* in human (56). Furthermore, the signaling strength of each interaction was predicted, as seen in the exSigNets plot (Figure 3G-H).

Cell differentiation often involves transient bifurcation between stable cell states (57), which may be analyzed using the concept of critical transition (CT) (Table 1). Previously computational methods have been developed to predict and quantify the critical transition and infer the transcription factors that regulate this transition through concepts such as critical transition signals (CTSs) (58-60). To investigate the capability of exFINDER in identifying CTS and its connections with external signals, we first used BioTIP (60) to predict CT using the index of criticality (Ic) and infer CTS from single-cell transcriptomes data. By direct usage of BioTIP, we computed the Ic of each NC cell group, and found that the transitional cells undergo a critical transition (Figure 3I), a result consistent with a previous finding (46). Next, we used exFINDER to infer the connections between the external signals to the top 10 marker genes of pigment cells and skeletal cells. Three external signals (*igf1*, *itgb1b2*, and *gdnfa*) are found to activate both pigment and skeletal cells through two CTSs (*tal1* and *lmo2*, inferred by BioTIP) (Figure 3J). This study further shows the capability of exFINDER in identifying external signals and their connections with critical transition during cell differentiation.

### exFINDER suggests critical external signals and targets during sensory neurogenesis in mouse

Using a mouse data on the early stages of sensory neurogenesis, we show how exFINDER can be utilized to predict the dominant signal sources and targets based on their signal inflows and outflows. The scRNA-seq data for mouse includes the somatosensory neuro-glial progeny of the trunk at E9.5, E10.5, E11.5, with 10 cell groups (Supplementary Figure S6A-B) (61). The original study inferred trajectories containing multiple cell fate decision points (Supplementary Figure S6C-E). One important fate choice-related trajectory is from unassigned cell group 3 (UA.3) to mechanoreception and proprioception cells (Figure 4A), leading to a fate choice between proprioceptor and mechanoreceptor lineages. Next, we study an unsolved question: what external signals the differentiating cells may receive and how they activate the cell differentiation? Since both mechanoreception and proprioception cells are differentiated from UA.3 cells, we first performed exFINDER analysis to infer the ligand-target GRN from the top 10 marker genes of the proprioception cells. exFINDER identified four external signals not expressed in all measured cells. However, their corresponding exSigNet shows five marker genes through highly expressed receptors and TFs in the data whereas these four receptors only interact with the external signals (Figure 4B-C). Interestingly, the interactions between *Pld2* and *Arf1*, *Slurp1* and *Chrna4* are supported by existing studies (62-64).

**Figure 4.**
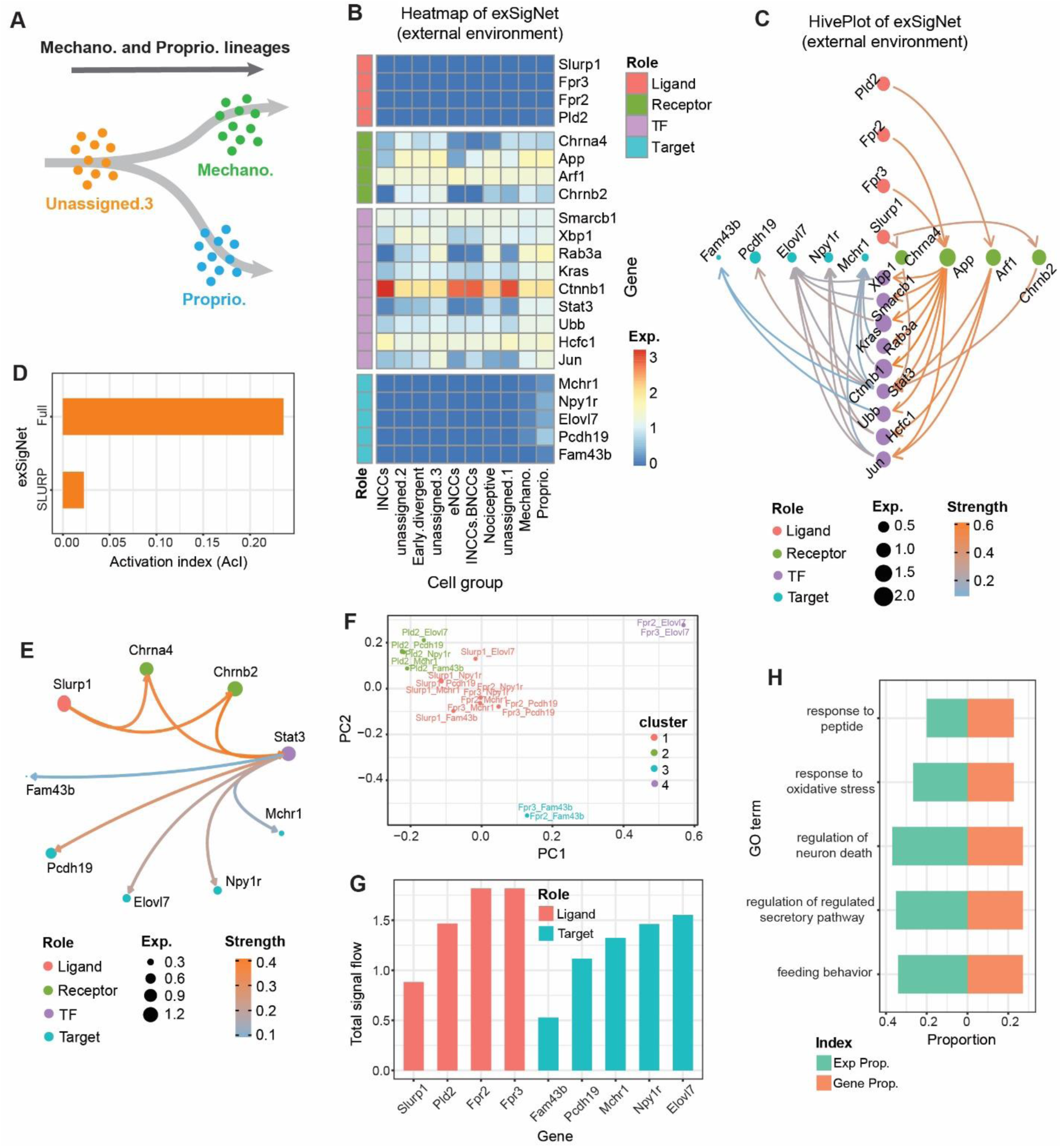
exFINDER identifies external signals and predicts critical signal sources and targets during sensory neurogenesis in the mouse. **(A)** Schematics of neural crest cell lineage showing cell differentiation from UA.3 cells to the mechanoreception and proprioception cells (61). **(B)** Expression heatmap of the exSigNet associated with the inferred external signals targeting the proprioception cell marker genes. **(C)** Hive plot showing the exSigNet. **(D)** Activation index of the full exSigNet and SLURP-related exSigNet. **(E)** Circle plot showing the SLURP-related exSigNet. **(F)** Clustering of the signaling network between different ligand-target pairs. **(G)** Bar plot of the total signal inflows and outflows of each external signal and target gene, respectively. **(H)** The expression and gene proportions of top 5 GO terms projecting to the exSigNet.

By comparing the ligand-receptor pairs of the exSigNet to a published ligand-receptor dataset (20), we found that part of the exSigNet (the activation from *Slurp1* to *Chrna4* and *Chrnb2*) is related to the SLURP pathway, a finding supported by previous studies (65, 66). Then we inferred the SLURP pathway-related part and compared its activation level to the exSigNet using the activation index (AcI). We found that although *Slurp1* activates two receptors, their overall activation levels are only moderately high (Figure 4D). This can be explained by the observation that *Chrna4* and *Chrnb2* only interacted with one TF out of nine (Figure 4E).

Since an exSigNet usually contains multiple signals and target genes, comparing the signaling networks between different ligand-target pairs is useful in uncovering underlying mechanisms. Thus, we classified the signaling networks between every ligand-target pair based on their structures and interaction strengths (Figure 4F). Four groups are found. All *Slurp1*-related pairs are grouped as one since they all belong to the SLURP pathway. *Pld2*-related pairs are in one group, since *Pld2* only activates *Arf1*. However, *Fpr2* and *Fpr3*-related pairs are mixed and clustered into three different groups, showing their potential roles for different biological events.

To quantify and compare the importance of external signals and target genes, we computed the maximal signal flow between every ligand-target pair and ranked the external signals and target genes based on their total signal outflows and inflows, respectively (Figure 4G). It is found that *Fpr2* and *Fpr3* have the most signal outflows – critical external signals (Table 1), while *Elovl7* has the most signal inflow – a critical target. *Slurp1* has the smallest signal outflow, which is consistent with our previous analysis that SLURP pathway-related signaling network does not have a high activation level.

exFINDER can also be used to evaluate the GO analysis outputs by projecting the top 5 GO terms (with the least *p*-values) to the exSigNet. Unlike GO analysis, which only uses gene symbols to infer the related biological events, exFINDER compares the GO term-related expression level and GO term-related genes to the exSigNet (Figure 4H), for example, showing that regulation of neuron death may be a critically important event in the exSigNet, although it has the third least p-value in GO analysis.

### exFINDER predicts the roles of external signals and uncovers transition paths in differentiation

During mouse sensory neurogenesis, besides the differentiation to mechanoreception and proprioception cells, several transition trajectories were also identified (61), including the one from early neural crust cells (eNCCs) to late neural crust cells (lNCCs) and then to boundary cap cells (INCCs/BCCs) (Supplementary Figure S6E), as well as two main neurogenic branches starting from INCCs and INCCs/BCCs, respectively (Figure 5A, Supplementary Figure S6C-D). These two branches both show the transition from progenitors to post-mitotic newborn neurons and may intersect through UA.1 cells (Figure 5A). To figure out which cell group undergoes the stronger transition in the multiple branches, we first performed BioTIP analysis that shows INCCs are most likely to be the transition cells (Figure 5B). This is reasonable because INCCs are in between of eNCCs and INCCs/BCCs, and the starting point of Branch A. Then exFINDER identifies that external signals (*Fpr2* and *Fpr3*) interact with one CTS (*Mitf*) by activating receptor *App*, and CTS (*Mitf*) activates the marker genes of both INCCs, INCCs/BCCs, and UA.1 cells (Figure 5C).

**Figure 5.**
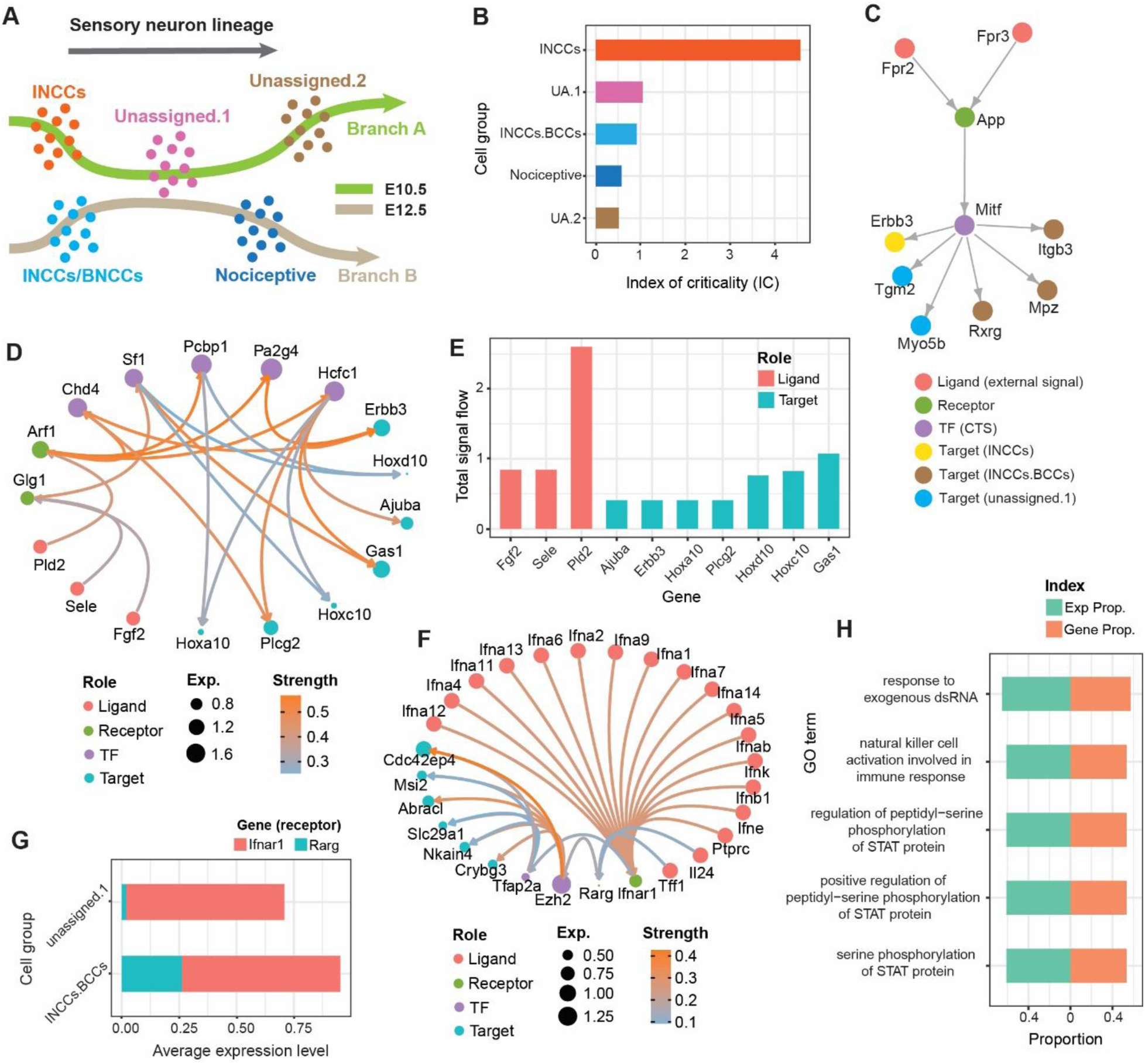
exFINDER suggests roles of external signals in different trajectories and predicts the transition paths during mouse sensory neurogenesis. **(A)** Schematics showing two branches and the corresponding cell groups. **(B)** Bar plot showing the index of criticality of cell groups along two branches generated by BioTIP. **(C)** The network inferred by exFINDER showing the critical transition signal-involved exSigNet. **(D)** Circle plot for the exSigNet linking external signals and marker genes of INCCs. **(E)** Bar plot of the total signal inflows and outflows of each external signal and target gene, respectively. **(F)** Circle plot showing the exSigNet linking external signals and marker genes of nociceptive cells. **(G)** Bar plot of the expression levels of the receptors in different cell groups. **(H)** The expression and gene proportions of top 5 GO terms projecting to the exSigNet.

Since INCCs are most likely to be the transitional cells, and CTS-involved signaling is likely not specific for INCCs, we next study how the external signals drive the expression of INCCs. In addition, signals from the external environment can be critical during the formation of INCCs as these cells are sampled early (E10.5). Then we employed exFINDER to identify external signals that targeted the top 10 marker genes of INCCs, showing that 7 out of the top 10 marker genes are activated by three external signals (Figure 5D). And all the ligand-receptor interactions have been confirmed in a previous study, such as *Pld2-Arf1* (67), *Fgf2-Glg1* (68), and *Sele-Glg1* (69). Meanwhile, exFINDER inferred a critical external signal *Pld2* and marker gene *Gas1* based on the signal flow (Figure 5E).

Along Branch A we observe a transition from INCCs to UA.1 cells and then UA.2 cells (Figure 5A), however, the transition from UA.1 cells to nociceptive cells may also be possible, suggesting the external signals that regulate INCCs may also interact with the nociceptive cells. Meanwhile, a group of early divergent genes along Branch B and their related TFs were identified in (61) (Supplementary Figure S6F). So, to investigate this hypothesis, we used exFINDER to identify the external signals that activate the divergent genes through their related TFs (Figure 5F). Interestingly, *Ifnar1* is the receptor of 12 subtype *Ifna* genes, while a previous study pointed out that the interferons act directly on nociceptors to produce pain sensitization (70). The comparable expression levels of two receptors (*Ifnar1* and *Rarg*) in UA.1 cells and INCCs/BCCs showing that *Rarg* has comparable expressions in both two cell groups, support our hypothesis (Figure 5G). These findings suggest multiple external signals are interacting with one receptor, which is highly expressed in both UA.1 cells and INCCs/BCCs, indicating a potential transition from INCCs to nociceptive cells. Last, we applied GO analysis and used exFINDER to evaluate the top five GO terms (Figure 5H). The results suggest the top five GO terms with the smallest p-values also have very similar expression values and gene proportions, indicating that they may have similar significances in the signaling from external signals to the early divergent genes.

### exFINDER uncovers the externally added inducers, revealing signaling pathways driving EMT

The epithelial-mesenchymal transition (EMT) is a critical cell fate transition. In a recent study that uncovered the context specificity of such process (34), a comparative analysis was performed for the EMT response with three different added inducers (*TGFB1*, *EGF*, and *TNF*) and scRNA-seq was used to measure expression profiles of four human datasets (A549, DU145, MCF7, and OVCA420). The study also compared 12 EMT time course experiments and investigated the effects of each inducer and the differential expression of the EMT regulators in all cases. For every time course dataset, we separated the cells into three groups based on the treatment (before, on, and off treatment), then applied exFINDER to all 12 datasets to infer the external signals targeting the EMT regulators.

In three *TGFB1*-induced datasets, *TGFB1* was found as one of the external signals and inferred the corresponding exSigNets (Figure 6A, D, and Supplementary Figure S7). In the *TGFB1*-induced MCF7 cells, the *TGFB1*-associated signaling network shows that the receptor *ITGB1* has a low expression, although its downstream transcription factors are still highly expressed (Supplementary Figure S7). Such differences are likely due to that these transcription factors are sensitive to the signal from *ITGB1*, or they are activated by other signals that are not associated with *ITGB1*. Meanwhile, in all four *TNF*-induced datasets, exFINDER successfully identified *TNF* as one of the external signals and inferred the corresponding exSigNets (Figure 6B, E, and Supplementary Figure S7), even if it was not measured in the data (Figure 6B). Our analysis shows *CALM1* and *TNFRSF1A* are two receptors that interact with *TNF*, a result supported by a recent study (71). In the four *EGF*-induced datasets, although its receptors are lowly expressed, the downstream transcription factors are still highly expressed (Figure 6C, Supplementary Figure S7). This indicates possible existence of other external signals that may activate those transcription factors. Especially in the A549 cells, *EGF* only regulated one target gene (Figure 6C), consistent with the fact that no significant increase in EMT score was observed in this case. As seen in the exSigNets, four receptors (*ITGB1*, *M6PR*, *CALM1*, and *TNFRSF1A*) were identified to interact with the inducers, and the comparison of their expression levels in different time course experiments indicates (Figure 6F): (1) *ITGB1 having* higher expression with the *TGFB1*-treatment in three out of four datasets; (2) No obvious change in expression levels for the *EGF*-associated receptor *M6PR* in the A549 cells; and (3) no expression increase in the *TNF*-associated receptors (*CALM1* and *TNFRSF1A*) in the OVCA420 cells. All the findings are consistent with the change in EMT scores shown in the previous study (34).

**Figure 6.**
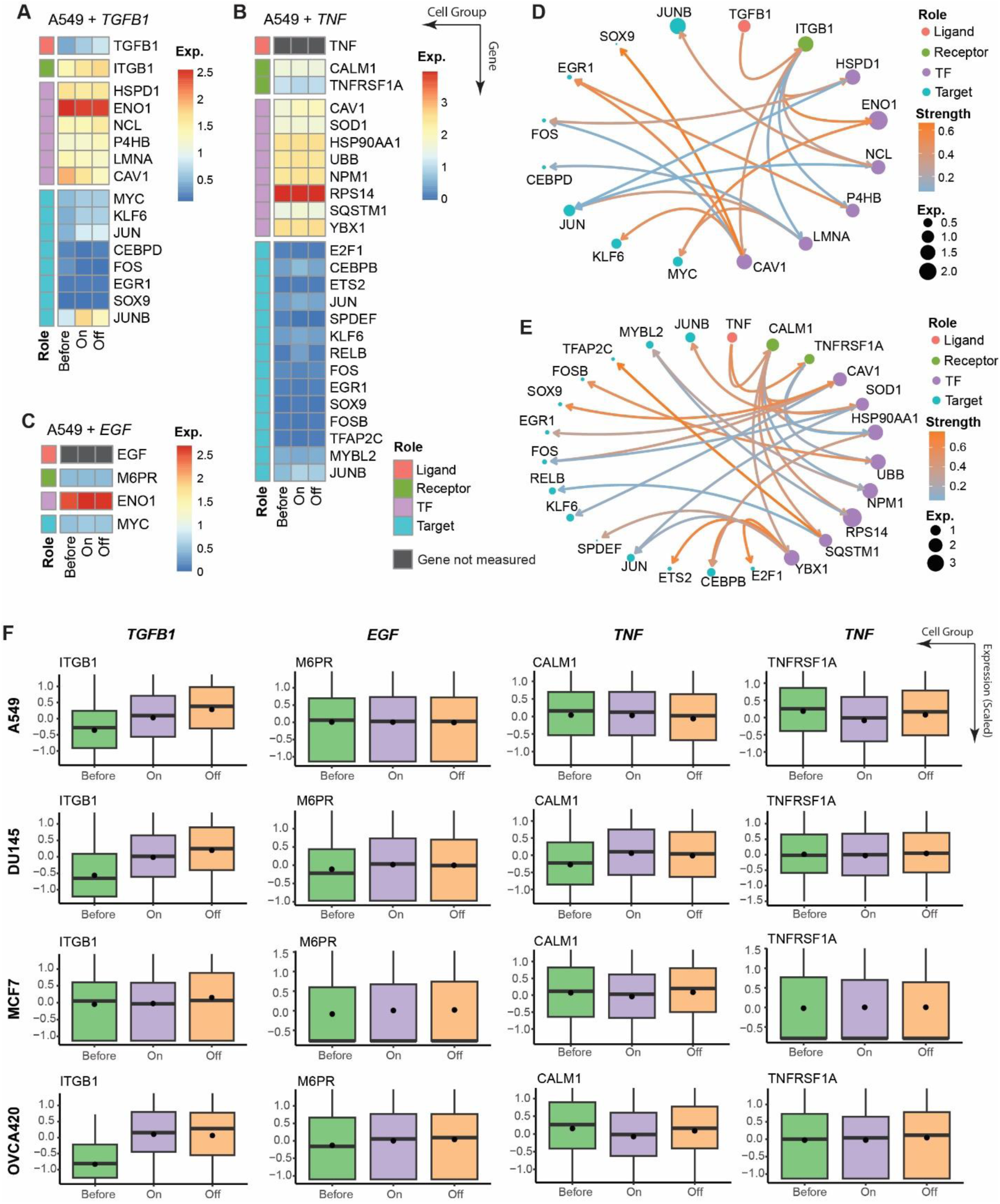
exFINDER uncovers the externally added inducers, revealing signaling pathways driving EMT. **(A-C)** Expression heatmaps of the exSigNets associated with the inferred external signals targeting the EMT regulators under different inducer-treatments in the A549 cells. **(D-E)** Circle plots showing the *TGFB1*-related and *TNF*-related exSigNets under the corresponding inducer-treatment in the A549 cells. **(F)** Box plots of the inferred receptors in different human cell types under different inducer-treatments, solid bar represents the median and the black dot represents the average.

## DISCUSSION

Here we have presented a new computational method to identify external signals in cell communication, infer their associated signaling networks and quantitatively analyze their downstream genes using scRNAseq data and prior knowledge. Central to this method is the exSigNet which contains: (1) signaling paths that link the external signals and target genes, and (2) the edge weight representing the predicted signaling strength. By applying exFINDER to recently published datasets, and with comparison to the other methods, our method was shown to be able to identify external signals consistent with the existing knowledge. It is worth noting that the utilization of exFINDER databases and scRNA-seq data provides cross-validation from both biological and statistical approaches, allowing robustness of the method. To our knowledge, exFINDER is the first computational method that can systematically identify external signals in cell communication, infer and quantitatively analyze the networks that connect the signals and downstream genes.

An R package has been developed with a user-friendly toolkit for inferring, analyzing, and visualizing external signals and the exSigNet based on any given scRNA-seq data. Different visualization outputs are provided, including customized circle plot and hive plot to present the structure of the exSigNet, the expression level of involved genes, and predicted signaling strength within the network. Meanwhile, exFINDER also allows easy exploration, download, and update of the built-in databases for different animal species. Any new progress in classifying cell groups and inferring trajectories (e.g., Monocle (10,72,73)) from scRNA-seq data can be easily included in the preprocessing steps to further improve accuracy and robustness of exFINDER. The flexibility of using different sub-modules and the interoperability in exFINDER allow users to take advantage of various utilizes in the package.

During the exFINDER analysis, the mass-action law, which has been widely used in estimating protein and mRNA activity levels (20, 74), plays an important role in quantifying the signaling strength and activation level. While the Hill function is a plausible approximation for modeling the nonlinearity of protein interactions, estimating the biologically reasonable ranges of the parameters (e.g., Hill function coefficient and dissociation constant) remains challenging.

The external signal inference for complex heterogeneous data remains a major challenge due to the lack of ground truth (14, 75). For some cases, because the cell lineage and cell population groups are not well studied, the inference of ligand-receptor interactions by cofactors, including agonist and antagonist, becomes difficult. One way for improvement is to include potential bidirectional interactions between certain genes when the ligand interaction structures (e.g., with multiple subunits).

Another possible improvement is to utilize the CITE-seq data, which contains the Antibody-Derived Tag (ADT) data for selected proteins, to identify the genes with ADT. To study this point, we analyzed a dataset of 8,617 cord blood mononuclear cells (CBMCs) produced with CITE-seq ((76), GSE100866) using exFINDER which contains the antibody-derived tag (ADT) data of 10 genes. A previous study has identified 15 cell population groups and their corresponding marker genes (https://broadinstitute.github.io/2020_scWorkshop/cite-seq.html, Supplementary Figure S8A). To perform exFINDER analysis, we first set the NK cells as the target cells and employed exFINDER to infer the external signals targeting its top 10 marker genes. Although six external signals (*PRL*, *PIK3CB*, *MDK*, *PTN*, *FPR2*, and *PLD2*) were identified, none of them was part of the 10 ADT-measured proteins (Supplementary Figure S8B-C). This is likely because the number of ADT-measured proteins is very small (only 10 proteins in this dataset), especially compared to the number of measured genes in RNA sequencing. In addition, we set the *CD14*^+^ Mono and *CD16*^+^ Mono cells to be the external cells and used exFINDER to infer the external signals expressed by them targeting the top 10 marker genes of NK cells. *CD34* was identified as the external signal targeting two marker genes *GZMB* and *PRF1* (Supplementary Figure S8D-E). However, according to the ADT data, *CD34* is highly expressed in several cell population groups including *CD14*^+^ Mono cells (Supplementary Figure S8F). These results suggest using CITE-seq with sufficient ADT data can further improve the inference accuracy.

In all, with the advances in different single-cell omics techniques and the development of computational methods, exFINDER may be combined with other data modularity and methods. For example, the directed dynamic information provided by RNA velocity (77) and cell transition paths inferred by MuTrans (78) can potentially be used in exFINDER to investigate the critical external signals that drive the cell fate transitions. The spatial omics technologies, such as spatial-CITE-seq (79), may be used to scrutinize the external signals coming from different spatial locations or directions, and the signaling strength based on expression levels can be finetuned by spatial distances between cells for more accurate inference constrained by spatial ranges of diffusive ligands. It is also useful to integrate more prior knowledge of spatial signaling into exFINDER-DB, enabling identification of the distance-dependent exSigNet using spatial transcriptomics data.

## AVAILABILITY

exFINDER is an open-source R package available in the GitHub repository (https://github.com/ChanghanGitHub/exFINDER.git)

## SUPPLEMENTARY DATA

Supplementary Data are available at NAR online.

## ACKNOWLEDGEMENT

We acknowledge helpful discussions with Suoqin Jin and members of the Thomas Schilling Lab. Author contributions: Q.N. designed the study, managed the project, and interpreted the results. C.H. developed the method, drafted the manuscript, built the online tutorial, created, and maintains the Github repository and the R package. P.Z. contributed to the R package and interpreted the results. All authors proofread and edited the manuscript.

## Supplementary Materials

**Figure S1.**
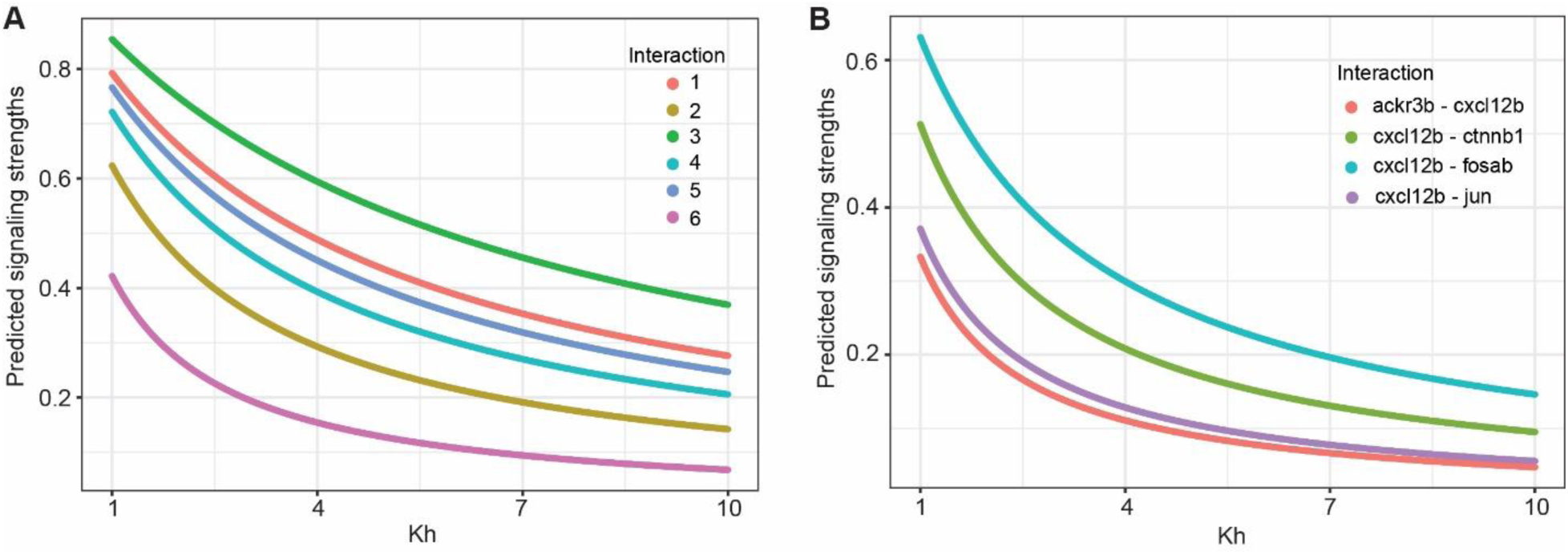
Predicted signaling strengths with different *Kh* values, showing that changing the value of *Kh* from 1 to 10 (with step size 0.01) does not significantly affect the relative signaling strengths (i.e. the order of signaling strength) of communication edges. **(A)** Dot plot showing the predicted signaling strengths of the synthetic data. Each interaction contains two genes with random expression values between 0 to 5. **(B)** Dot plot showing the predicted signaling strengths of four zebrafish gene interactions. The corresponding gene expression data is obtained from Tatarakis et al. 2021.

**Figure S2.**
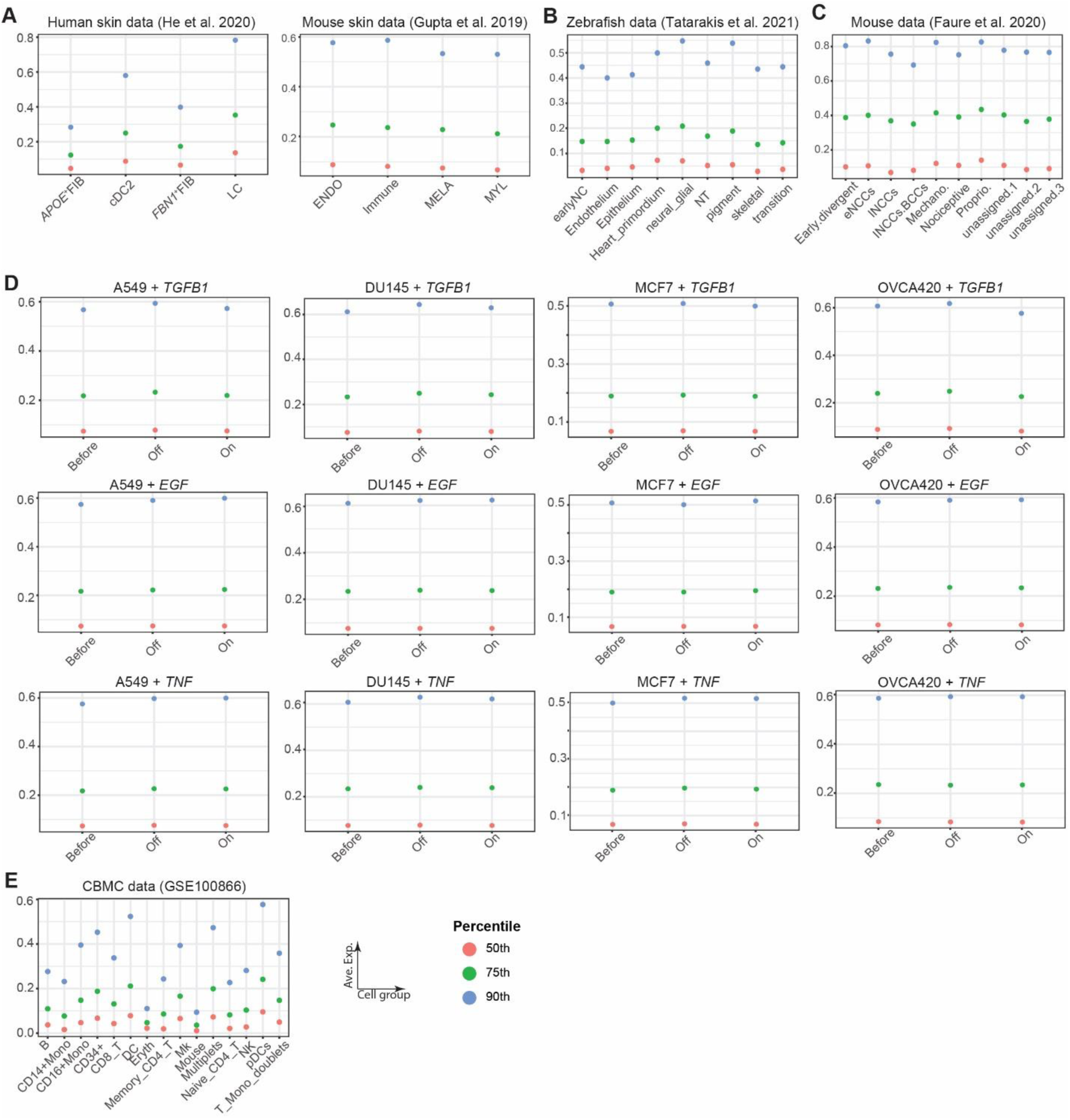
Percentiles for determining cutoff levels for “lowly expressed” and “highly expressed” genes in datasets used. 50^th^, 75^th^ and 90^th^ percentiles of **(A)** Human skin data (He et al. 2020) and Mouse skin data (Gupta et al. 2019), **(B)** Zebrafish data (Tatarakis et al., 2019), **(C)** Mouse data (Faure et al. 2020), **(D)** A549, DU145, MCF, OVCA420 data (Cook et al. 2020), and **(E)** cord blood mononuclear cells (CBMCs) data (Stoeckius et al. 2017, GSE100866).

**Figure S3.**
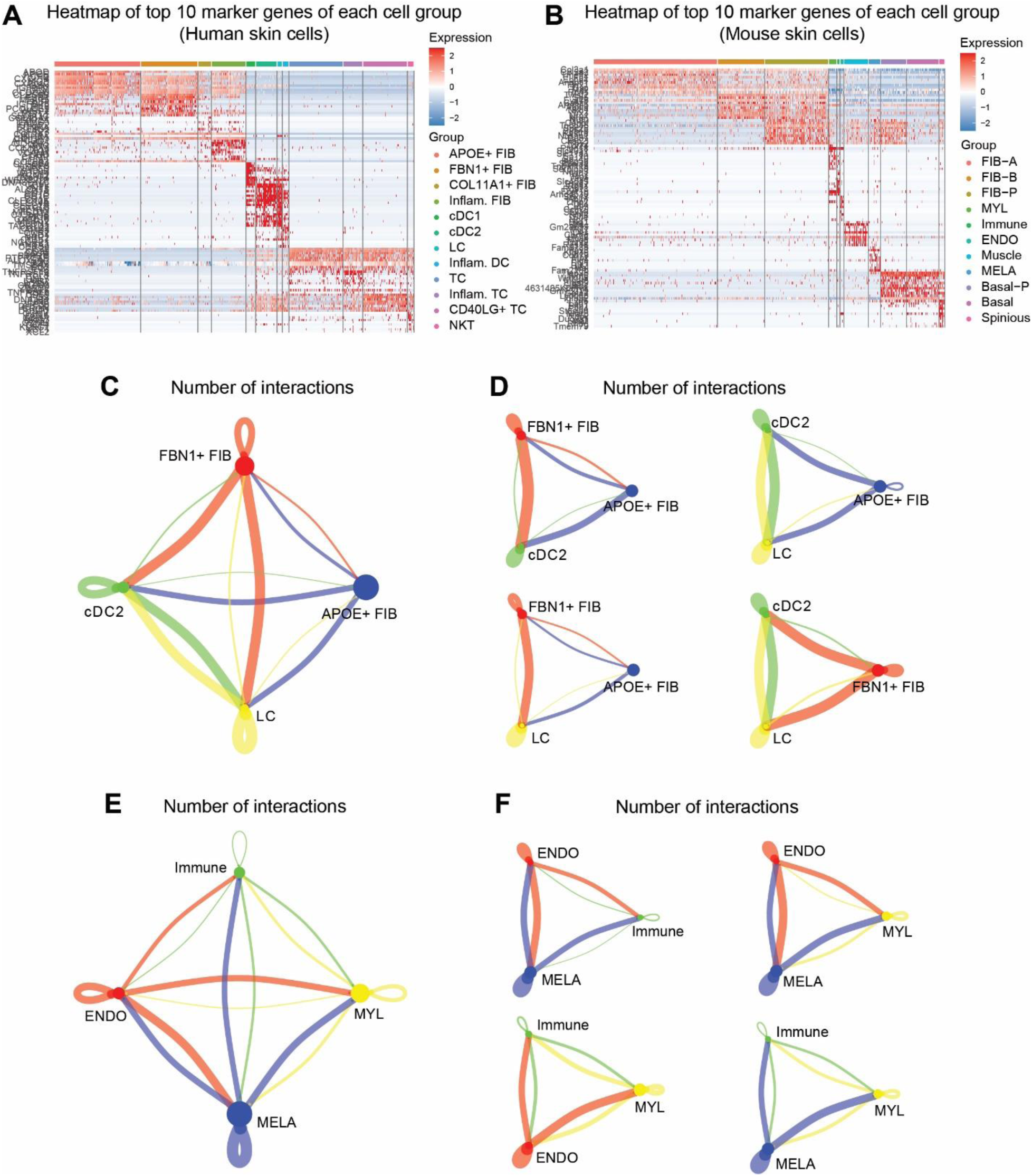
Marker gene expressions and CellChat inferred cell-cell communications in skin datasets. **(A-B)** Heatmap of top 10 marker genes of each human skin and mouse skin cell group (generated by Seurat). **(C-D)** and mouse skin cells **(E-F)** inferred by CellChat. **C, E** Number of interactions between four cell groups. **(D-F)** Number of interactions between every three cell groups.

**Figure S4.**
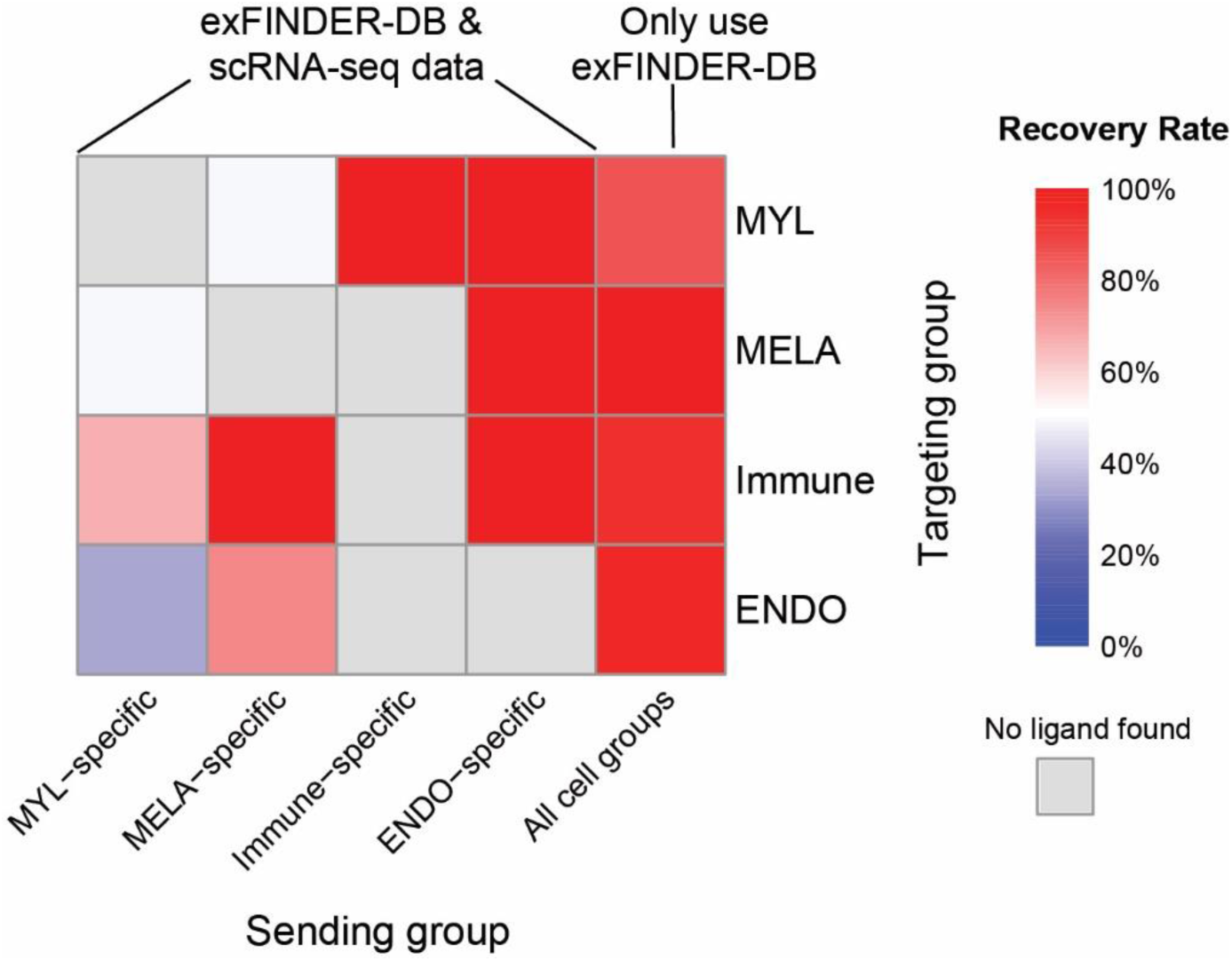
Benchmarking results of exFINDER using dataset of mouse skin cells. Heatmap showing the percentages of ligands inferred by CellChat that also captured by exFINDER. The X-axis represents the cell population groups expressing the ligands, and Y-axis represents the cell population groups receiving the signals. The color bar is the percentages of CellChat-inferred ligands that are also identified by exFINDER. The 5^th^ column shows the ligand recovery rate of exFINDER only using prior knowledge, the rest four columns show the ligand recovery rate of exFINDER using both prior knowledge and the scRNA-seq data. And the grey block indicates no such ligands inferred by CellChat.

**Figure S5.**
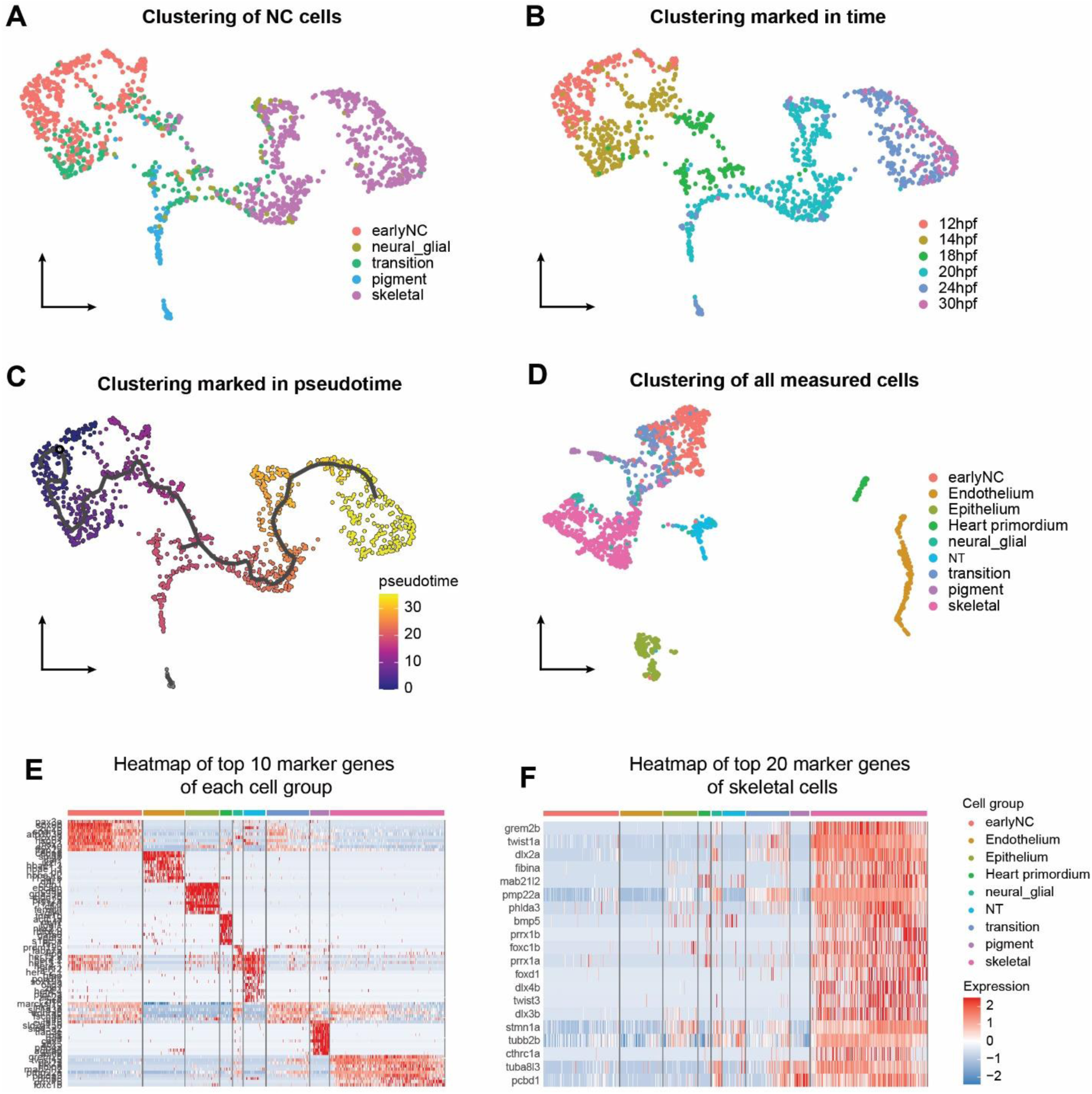
Overview of the dataset of zebrafish neural crest (NC) cells and marker gene expressions (generated by Seurat). **(A)** Reproduction of the dimensionality reduction results of the scRNA-seq data using Seurat. **(B)** Visualization of the dimensionality reduction with time information. **(C)** Trajectory analysis and visualization with pseudotime values using Monocle3. **(D)** Reproduction of the dimensionality reduction results of all measured cells, including NC cells and non-NC cells. **(E)** Heatmap of top 10 marker genes of each cell group. **(F)** Heatmap of top 20 marker genes of skeletal cells.

**Figure S6.**
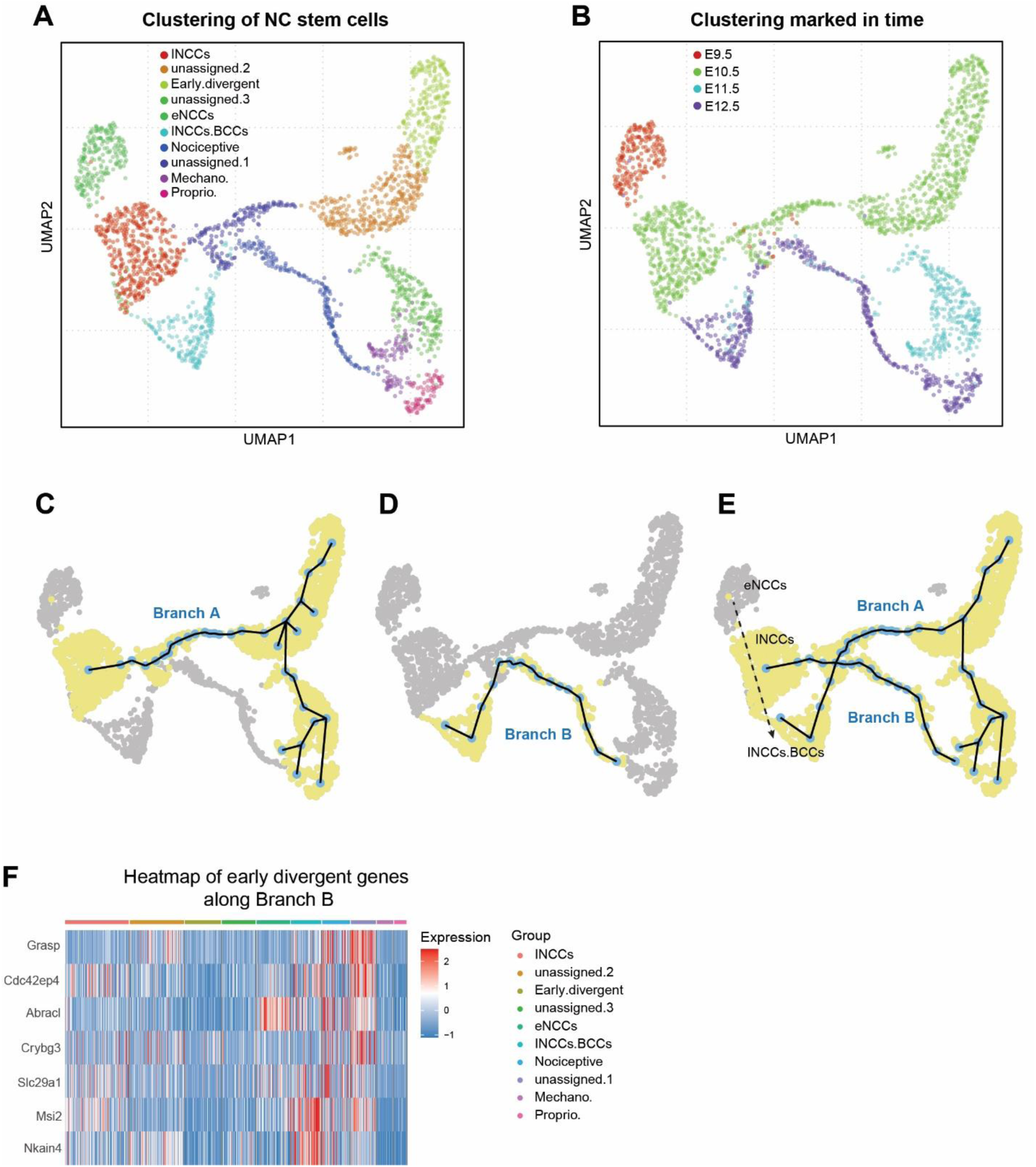
Overview of the dataset of mouse neural crest (NC) cells and marker gene expressions (generated by Seurat). **(A)** Reproduction of the dimensionality reduction results of the scRNA-seq data using pagoda2. **(B)** Visualization of the dimensionality reduction with time information. **(C-E)** Reproduction of the trajectory analysis results using provided codes. **(F)** Heatmap of the early divergent genes along Branch B.

**Figure S7.**
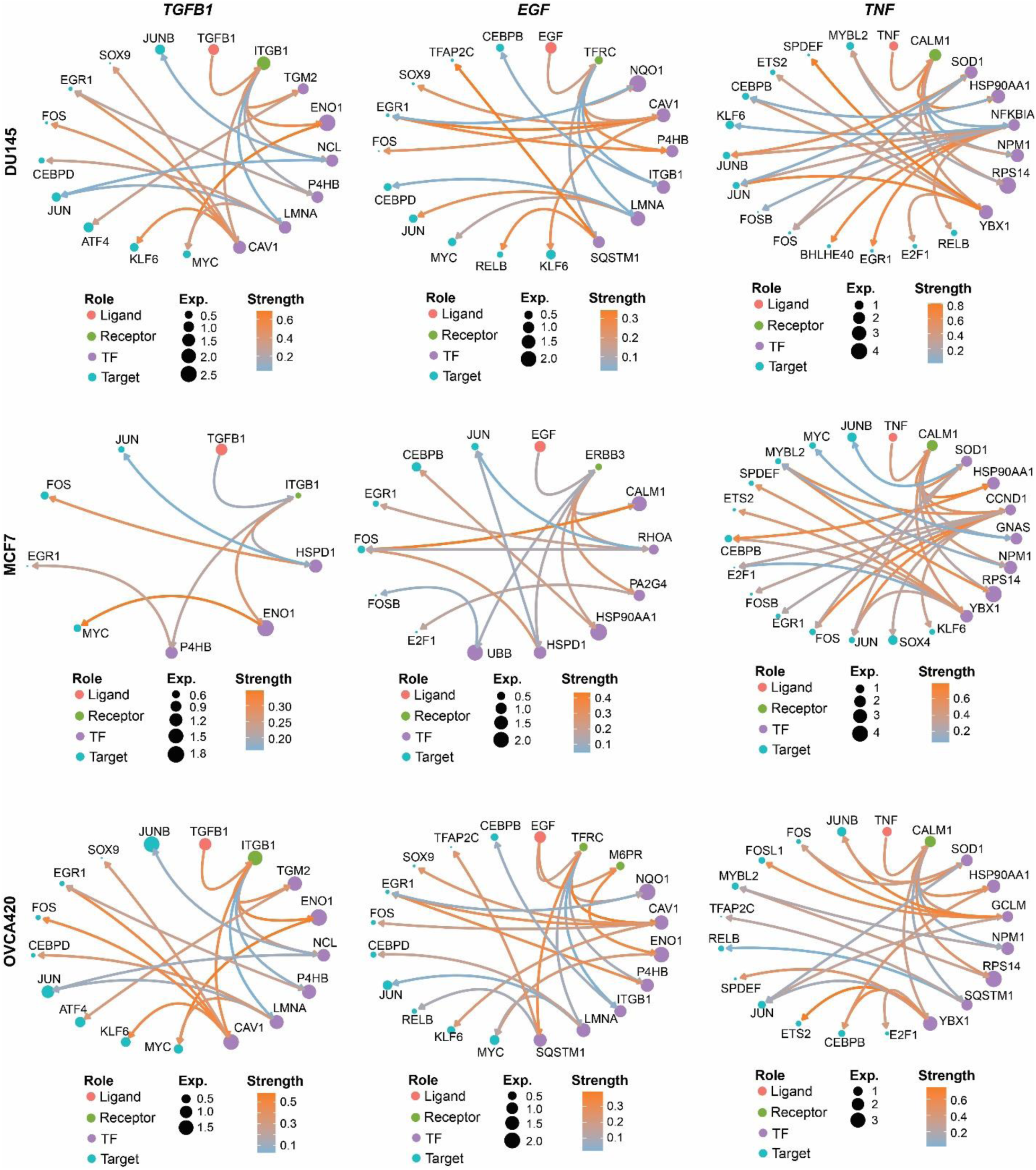
exFINDER inferred inducer-associated exSigNets in EMT datasets. Circle plots showing the inducer-associated exSigNets in DU145, MCF7, and OVCA420 cells.

**Figure S8.**
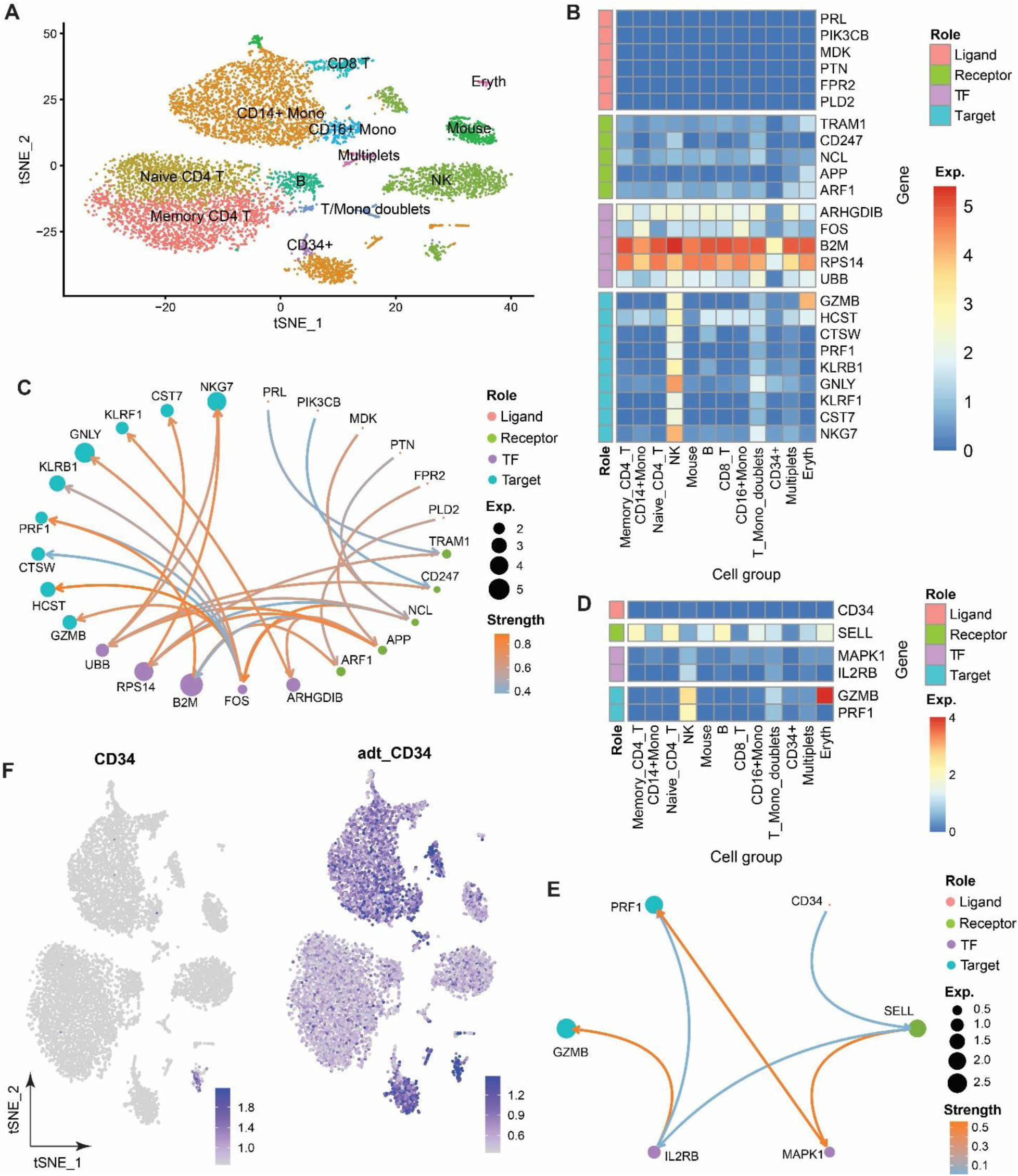
exFINDER analysis using the CITE-seq data of cord blood mononuclear cells (CBMCs). **(A)** Reproduction of the dimensionality reduction results of the CBMCs data using Seurat. **(B-C)** Expression levels and the circle plot of the exSigNet associated with the inferred signals came from the external environment targeting NK cell marker genes. **(D-E)** Expression levels and the circle plot of the exSigNet associated with *CD34* (which did not come from the *CD14*^+^ Mono and *CD16*^+^ Mono cells) targeting NK cell marker genes. **(F)** RNA expression level of *CD34* and its ADT level.

**Table S1.** The number of ligand-receptor, receptor-TF, and TF-target interactions recorded in four different databases, and their availabilities in different species. The number of interactions in a database is calculated based on the interactions with different genes associated with the sending and receiving signals. This quantity is subject to change when a database is updated.

|  | Number of interactions recorded (human) |  |  | Species |
| --- | --- | --- | --- | --- |
|  | ligand-receptor | receptor-TF | TF-target |  |
| exFINDER-DB | 13962 | 2475457 | 2963964 | Human, Mouse, Zebrafish |
| NicheNet | 12019 | 2475457 | 2912482 | Human |
| CellChatDB | 2021 | None | None | Human, Mouse, Zebrafish |
| ICELLNET | 1034 | None | None | Human |

